# Biophysical mechanism of ultrafast helical twisting contraction in the giant unicellular ciliate Spirostomum ambiguum

**DOI:** 10.1101/854836

**Authors:** L. X. Xu, M. S. Bhamla

## Abstract

The biophysical mechanism of cytoskeletal structures has been fundamental to understanding of cellular dynamics. Here, we present a mechanism for the ultrafast contraction exhibited by the unicellular ciliate *Spirostomum ambiguum*. Powered by a Ca^2+^ binding myoneme mesh architecture, Spirostomum is able to twist its two ends in the same direction and fully contract to 75% of its body length within five milliseconds, followed by a slow elongation mechanism driven by the uncoiling of the microtubules. To elucidate the principles of this rapid contraction and slow elongation cycle, we used high-speed imaging to examine the same-direction coiling of the two ends of the cell and immunofluorescence techniques to visualize and quantify the structural changes in the myoneme mesh, microtubule arrays, and the cell membrane. Lastly, we provide support for our hypotheses using a simple physical model that captures key features of Spirostomum’s ultrafast twisting contraction.

**SIGNIFICANCE:** Ultrafast movements are ubiquitous in nature, and some of the most fascinating ultrafast biophysical systems are found on the cellular level. Quantitative studies and models are key to understand the biophysics of these fast movements. In this work, we study Spirostomum’s ultrafast contraction by using high-speed imaging, labeling relevant cytoskeletal structures, and building a physical model to provide a biophysical mechanism especially of the helical same direction twisting of this extremely large single cell organism. Deeper understanding of how single cells can execute extreme shape changes hold potential for advancing basic cell biophysics and also inspire new cellular inspired actuators for engineering applications.

## INTRODUCTION

Ultrafast motility occurs at many length scales in nature, from large multicellular organisms such as a mantis shrimp and trap-jaw ants [1, 2] to microscopic single cells such as *Vorticella* and nematocysts of jellyfish [3–7]. Mechanistic understanding how organisms achieve repeatable rapid motion, especially at small scales is an emerging area of interdisciplinary research with potential to advance our understanding of the biophysics of cellular actuators and motors, as well as inspire design of small-scale robots [8]. In this work, our goal is to develop a biophysical understanding of the ultrafast contraction of the unicellular protozoan *Spirostomum ambiguum*, which upon an a stimulus can contract its 4 mm long cigar-shaped body to 1/4^*th*^ of its original length in less than 5 milliseconds, followed by a slow elongation mechanism (~1 s) back to its elongated length (Fig 1a,b, SI Movie 1) [9–14]. The duration of the contraction phase is so brief that 50 of these can fit in the blink of an eye [15]. The peak acceleration this unicellular organism achieves during its contraction is (*a*_max_ = 140 m/s^2^), which is ten times faster than that of a sprinting cheetah [16, 17]. Recently it was discovered that Spirostomum harnesses these ultrafast contractions to generate hydrodynamic trigger waves for intercellular communication and toxin dispersal in large cellular collectives [16].

**FIG. 1.**
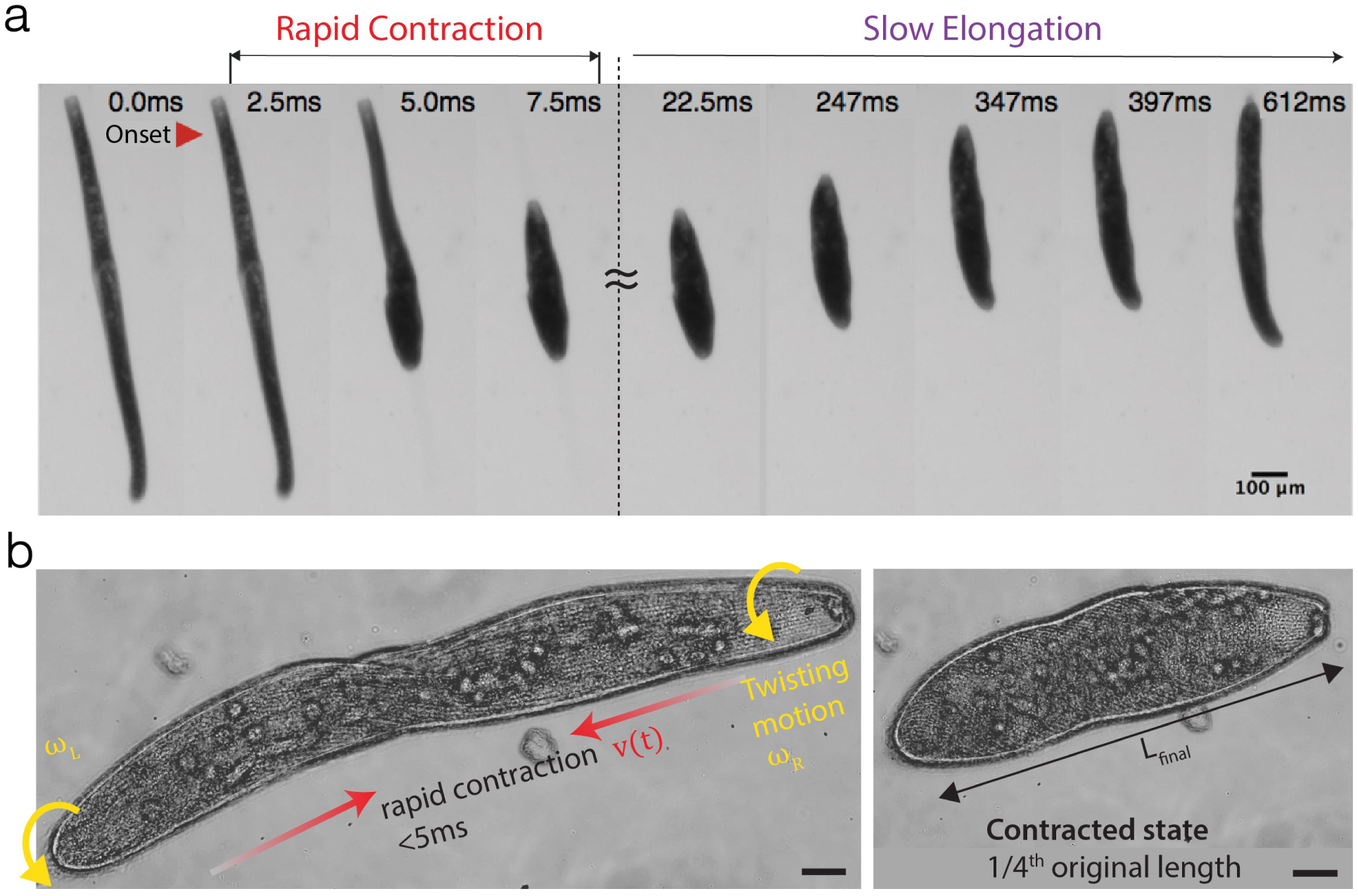
*Spirostomum ambiguum* contraction using high-speed microscopy. **a** Collage from video highlighting the rapid contraction and slow elongation of Spirostomum (SI Movie 1). **b** Same-direction twisting contraction in Spirostomum revealed by high-speed microscopy (20x) video at 2800 fps (SI Movie 2).

We focus on Spirostomum for two main reasons. First, due to its large size (4mm in length), Spirostomum is one of the largest free-swimming single cell organisms (readily visible by the naked eye). Thus, compared to Vorticella, another well-studied fast contracting protist, Spirostomum is 25*X* larger, but still produces 3X higher *v*_max_ and *a*_max_, offering a unique perspective into the physical limits of cytoskeletal components to generate power outputs over millimeter lengthscales (Table 1). Second, Spirostomum can repeatedly execute multiple contraction-elongation cycles during its life. Thus, unlike the one-shot destructive firing of nematocysts in cnidarians from high-pressure capsules, the governing mechanics of Spirostomum’s repeatable contraction offers insight into guiding principles for design of mechanically robust and reconfigurable cytoskeletal architectures.

**TABLE I.** Table comparing kinematics of three different unicellular ultrafast systems.

| System | $L$ ($\mu\text{m}$) | $\Delta L$ ($\mu\text{m}$) | $v_{\text{max}}$ ($\text{ms}^{-1}$) | $a_{\text{max}}$ (g) | $t$ (ms) | Motor | Repeatability | Ref |
| --- | --- | --- | --- | --- | --- | --- | --- | --- |
| *Spirostomum ambiguum* | 4000 | 3000 (75%) | 0.22 | 14 | 4.6 | Myoneme ($\text{Ca}^{2+}$) | rep | [16] |
| *Vorticella convallaria* | 180 | 117 (65%) | 0.072 | 10 | 6 | Spasmoneme ($\text{Ca}^{2+}$) | rep | [3–5] |
| Nematocyst | 14 | 4 (29%) | 28 | $5.4 \times 10^6$ | $7 \times 10^{-4}$ | Cnidoins (Osmotic stress) | one-shot | [6, 7] |

The ultrafast helical twisting contraction of Spirostomum is driven by a supramolecular assembly of myonemes fibers that comprise of a *Ca*^2+^-binding protein called centrin with potentially a hypothesized Sf1p-like backbone [18–23]. Using a high-speed camera mounted on a microscope, this twisting contraction can be appreciated in high resolution as shown in Fig. 1b. We note that unlike a piece of elastic string that can be contracted by twisting both ends in opposite directions, each end of the Spirostomum cell puzzlingly rotates in the same direction during contraction (Fig. 1b, SI Movie 2). To the best of our knowledge, a physical mechanism of this same direction rotational-contraction remains elusive. It is worth noting that contraction is not symmetric - there is a short time delay between the start of the motion of each end due to the propagation of a Ca^2+^ wave that travels along the cell body [4, 5]. After the contraction, Spirostomum slowly returns to its original shape by elongation.

Past work from the 70’s and 80’s utilized mainly electron and light microscopy on cross-sections of Spirostomum’s long body to understand the governing cytoskeletal components responsible for contraction [13, 14, 24]. Yagiu and Shigenaka [13] were the first to identify the existence of a contractile fibrillar system (myonemes). Lehman and Rebhun [24] proposed that myonemes may contract like muscles and result in the reconfiguration of microtubules, while the area of plasma membrane remained constant during cell body contraction. Hawkes and Holberton [25–27] rigorously examined Spirostomum’s contraction and relaxation kinetics in detail and identified myonemes as the mechanochemical engine that was powered by the *Ca*^2+^ ions. Yogosawa-Ohara, Suzaki, and Shigenaka [28, 29] observed that the myoneme fiber bundles (composed of 3-10 nm myoneme fibers) thickened during contraction, indicating that the myonemes were responsible for the shortening of the cell. They also hypothesized that the stiff microtubules were responsible for the helical twisting action during contractions. Their study was the first to conclude that the antagonistic action of the myonemes and the microtubules resulted in the repeatable contraction-elongation kinetics of Spirostomum [28, 29]. Ishida, Suzaki, and Shigenaka [30] built on Yogosawa-Ohara et al.’s model and proposed a physical linkage between the myonemes and the microtubules in order to transmit forces without slipping past each other.

Despite this impressive foundational body of work, a complete biophysical picture of Spirostomum’s reversible, and same-direction twisting contraction remains missing. Using modern high-resolution fluorescent imaging, here we aim to provide a quantitative analysis of the geometric rearrangements of the myonemes, microtubules and cell plasma membrane during contraction and elongation. We use these insights to provide a biophysical mechanism of Spirostomum’s ultrafast twisting contraction, and support our findings with a simple physical model.

## MATERIALS AND METHODS

### Spirostomum Culture

*Spirostomum ambiguum* were used for all experiments were obtained from Siento, UK and has been cultured in the laboratory. New cultures were prepared at 21°C. Culturing medium was prepared by mixing one part boiled hay medium solution (Ward’s Science, 470177-390), four parts boiled spring water, and two boiled wheat grains (Carolina, Item #132425).

### Fixation & Immunofluorescence

Cells were transferred from cultures into a 96-well plate and washed with spring water twice. To yield elongated samples, cells were permeabilized in 200 *µ*L of 0.5X PHEM buffer at 4°C, incubated for 6 hours and fixed with 100 *µ*L of 1X PHEM 2% PFA at 4°C. To yield contracted samples, cells were permeabilized in 100 *µ*L of 1X PHEM buffer at room temperature (RT) for 15 seconds, and fixed with 100 *µ*L of 1X PHEM 2% PFA for 15 minutes. Fixed cells were then treated with 300 *µ*L of 2% PFA 1X PHEM for 60 minutes, and washed three times with 300 *µ*L of 1X PBS 3% BSA 1% Saponin for 10 minutes each. To visualize cytoskeletal components following primary antibodies were used: myoneme; anti-Centrin (Millipore Sigma, Cat. #04-1624); microtubules; TAP952 (Millipore Sigma, Cat. #MABS277). Cells were incubated in 200 *µ*L primary antibody solution diluted to 10 *µ*L/mL in 1X PBS 3% BSA 1% Saponin for 15 hours at RT, and washed three times with 300 *µ*L of 1X PBS 3% BSA 1% Saponin for 10 minutes each. AF594 anti-Mouse IgG (ThermoFisher, Cat. #R37115) was used as the the secondary antibody. Cells were incubated in 200 *µ*L of its working solution in 1X PBS 3% BSA 1% Saponin for three hours at RT, and washed three times with 300 *µ*L of 1X PBS 3% BSA 1% Saponin for 10 minutes each. To stain the cell membrane, cells were incubated in 200 *µ*L CellMask Orange Plasma membrane Stain (ThermoFisher, Prod #C10045) diluted to 1 *µ*L/mL in 1X PBS 3% BSA 1% Saponin for 15 minutes and washed three times with 300 *µ*L of 1X PBS 3% BSA 1% Saponin for 10 minutes each.

For microscopy, stained cells were transferred onto a confocal coverslip (Neuvitro, Cat. #: GG-22-1.5-PLL) with 100 *µ*m spacers (Cospheric, CPMS-0.96 106-125 *µ*m). A glass slide (VWR, Cat. # 16004-430) were placed on the coverslip and inverted. To prevent evaporation, the slide was sealed with nail polish (FisherScientific, Cat. #72180) before observation in a Zeiss LSM 780 / Elyra PS1 Superresolution Microscope.

## RESULTS AND DISCUSSION

### Mechanism of Centrin-based Contraction

How do supramolecular assemblies of myoneme proteins generate contractile force over the millimeter length of the cell? To answer this, we first examine the structural changes in the myoneme mesh. Using anti-centrin labeling, we visualize the myoneme mesh architecture in both contracted and elongated states, revealing packed parallelogram-like microstructures that decorate the entire cell surface (Fig 2a). Bundles of myoneme filaments form the edges of each parallelogram, which consequently shorten to decrease the length of each side (*A*,*B*) of the parallelogram to generate an integrated contractile force *F* (Fig 2b). Indeed our data (Fig 2c) reveals a decrease from 3.8±0.8 *µ*m to 2.7±0.6 *µ*m in *A* (29%) and 4.5±1.2 *µ*m to 3.6±0.8 *µ*m in *B* (20%). The shortening of the fiber bundles is associated with a thickening of the filament bundles from 0.7±0.1 *µ*m to 0.8±0.12 *µ*m in *A* (14%) and from 0.6±0.1 *µ*m to 0.9±0.2 *µ*m in *B* (50%). The thickening of the myoneme bundles is in agreement with previous observations by Yogosawa-Ohara and Shigenaka [28, 29]. The myoneme bundle shortening consequently results in shearing of each individual parallelogram with an increase in the vertex angle *θ_AB_* from 54.4° ± 10.0° to 71.6° ± 9.5° (32%).

**FIG. 2.**
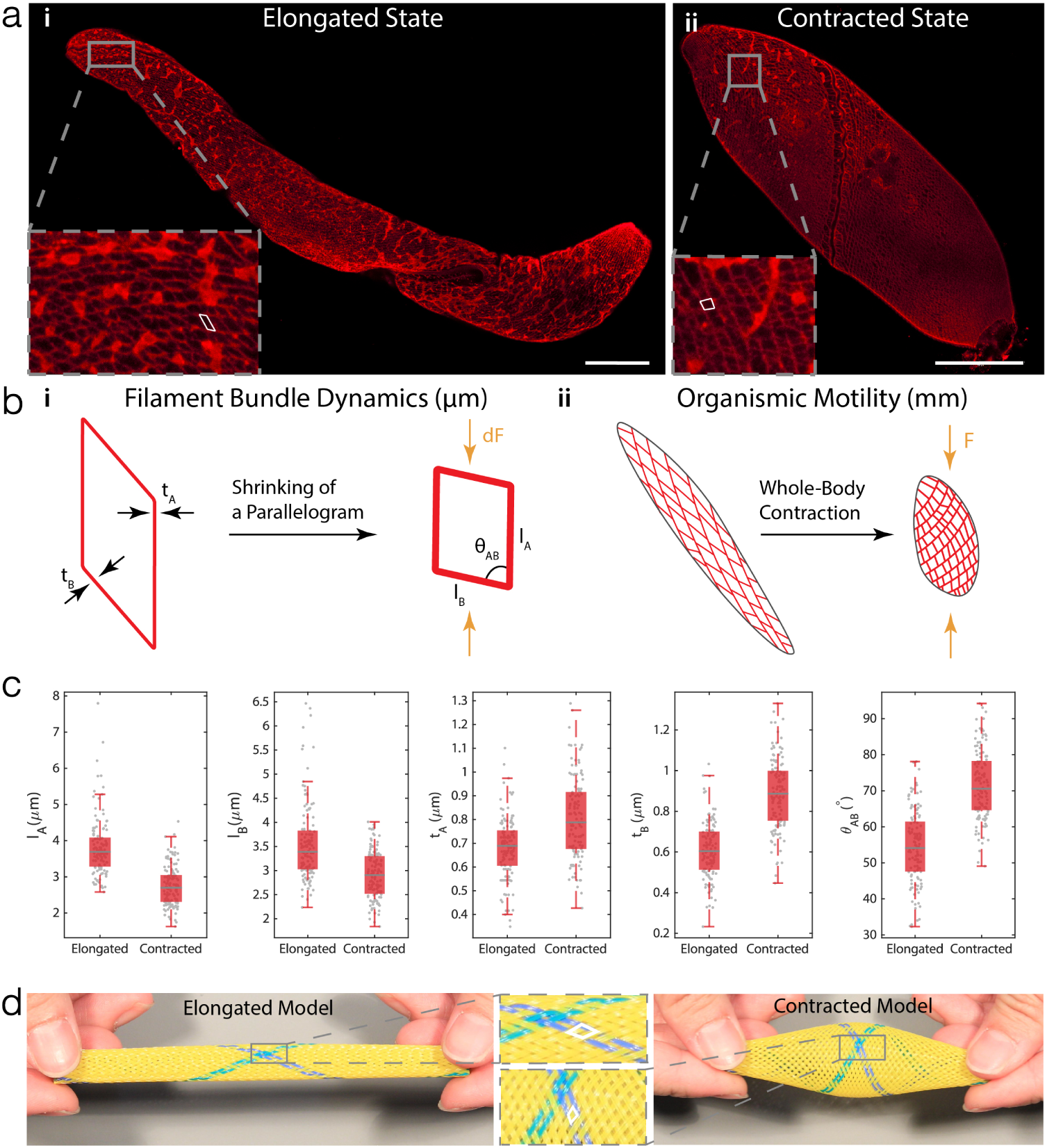
Mechanism of Centrin-based Contraction. **a** Immunofluorescence labelling of myoneme in both (i) elongated and (ii) contracted states. The myoneme mesh is comprised of individual “parallelogram” units (outlined in white). Scale Bar = 100 *µ*m. **b** A structural hierarchy of length scales (*µ*m - cm). (i) On the *µ*m-scale, each “parallelogram” shrinks as *Ca*^2+^ waves travel through the cell body. Such shrinking is quantified by the change in length of two sides (*l*_*A*_ & *l*_*B*_ , where *A* denotes the side that runs along the microtubules), the change in thickness of the sides *A* & *B* (*t*_*A*_ & *t*_*B*_ ), and the change in angle *θ*_*AB*_ between *A* & *B*. (ii) Myoneme mesh over the entire organism. Shrinking of individual parallelograms propagates along the cell body, causing the organism to not only shorten in length, but also change in angle *θ*_*AB*_ to cause a twisting motion. **c** Box-plots of quantitative measurements in the shape change of parallelograms (Elongated: N = 3 cells, 120 measurements for each box plot. Contracted: N =3 cells, 132 measurements for each box plot). As the cell contracts, myoneme parallelograms become shorter, and thicker. Gray lines in the middle of the boxes represent the median of the data set; the edges of the box represent the quartiles where the bottoms represent 25% and the tops represent 75% percentiles. The whiskers extend to the most extreme non-outlier data points. **d** A physical model of the myoneme mesh in its elongated and contracted states. The shape change of parallelogram units before and after contraction is similar to that of the actual cell.

Through quantitative analysis of these images, we can now propose a multiscale (*µ*m - mm) hierarchical physical mechanism of force-generation in myonemes to enable a full-body contraction. Bundles of these nanoscale myoneme fibers form the edges of the myoneme fishnet-like structure, which during contraction generate local contractile forces (*dF*) [28]. Since all the parallelograms are connected along the length of the body, the net contractile force is amplified (*F* ⋅ *N dF* ), where *N* denotes the number of parallelograms, resulting in a full-body ultrafast contraction.

To provide qualitative support for this proposed mechanism, we physically realize a fish-net structure to form a cylindrical shape mimicking Spirostomum’s cigar-shaped body. By compressing from both ends using ones fingers, and releasing, the myoneme physical analog follows the similar shearing of the individual parallelogram units (Fig. 2d, SI Movie 3). The fibers in this physical toy model are inelastic requiring manual force to compress them, while the actual myonemes behave as an entropic spring to self-generate contractile forces. We note that our proposed mechanism although novel in its description for myonemes in single cell organism, is a universal elastic network motif found also in multicellular worms (using collagen fibers) [31, 32] and engineered McKibben actuators in soft robotics [33].

### Reset and Elongation Mechanism due to the Microtubules

How does the the microtubule system serve as a reset mechanism to re-elongate the cell? Using confocal images of anti-tubulin stained cells (Fig 3a), we observe interestingly that the microtubules are arranged as transverse helices along the long axis of the cell body (Fig 3b). By measuring the average orientation of the microtubules with respect to the short axis of the cell (*θ*), we find that the contracted cells have smaller angles of *θ_C_* = 35.5° ± 9.5°(Fig 3c). From an energetic point, the microtubules are in an unfavourable higher bending energy configuration, since the bending energy of a transversely wrapped microtubule *E* ∝ cos^4^ *θ* [34]. From this expression, we note that the highest energy mode would be for smaller helical angles *E*_max_ = *E*(*θ* = 0), while energy is relieved as microtubules straighten out *E*_min_ = *E*(*θ* = *π/*2). Indeed as the cell elongates, the microtubules increase pitch angles, driving the cell in a lower energy state with microtubules almost parallel to the cell body *θ_E_* = 66.9° ± 12.2°.

**FIG. 3.**
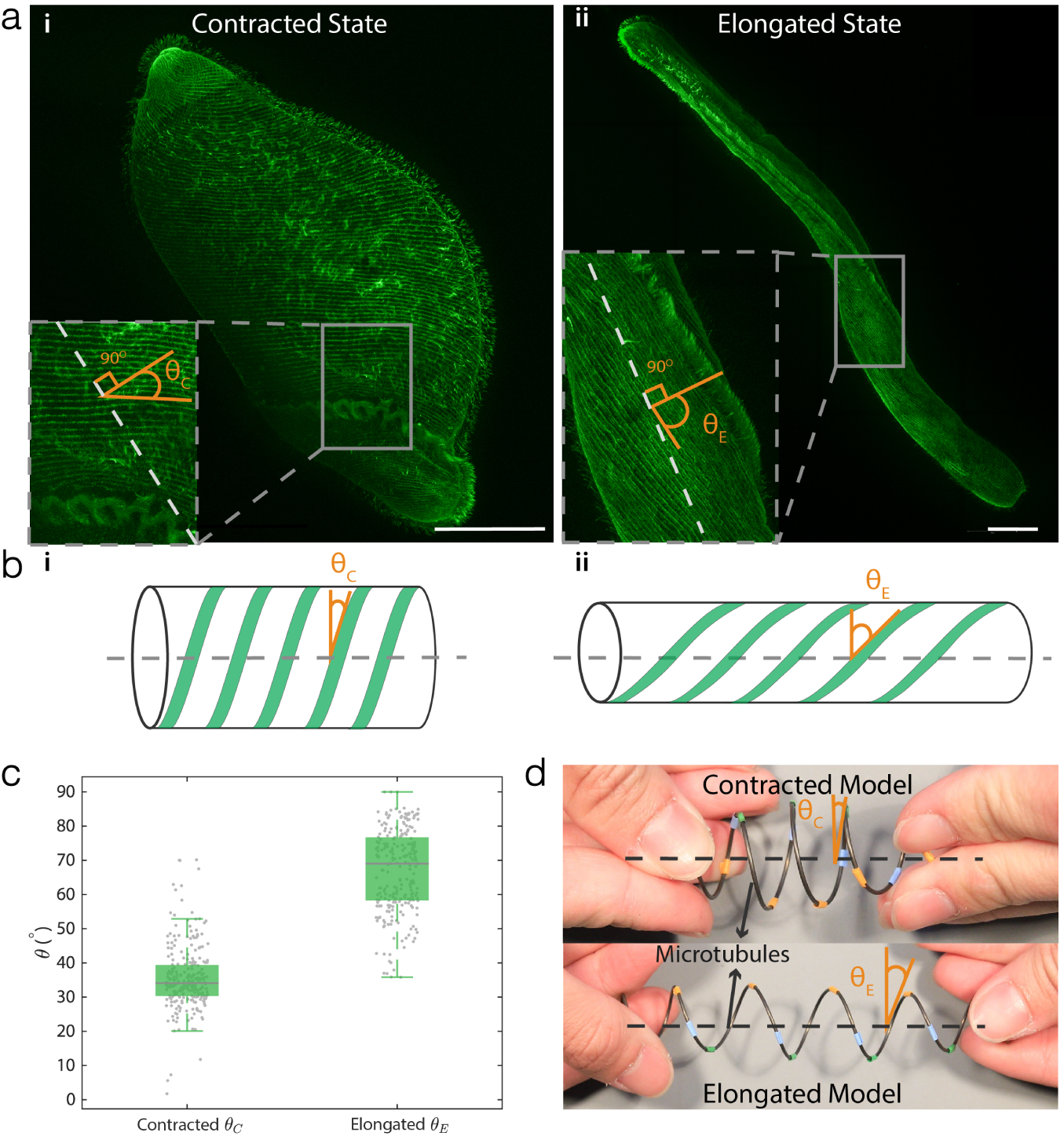
Reset and Elongation Mechanism due to the Microtubules. **a** Immunofluorescence labelling of microtubules in both (i) contracted and (ii) elongated state, with zoomed-in sections showing the angles of the microtubules with respect to the minor axis of the cell. Scale Bar = 100 *µ*m. **b** Illustration of changes in microtubule angles during the contraction in (i) contracted (*θ*_*C*_) and (ii) elongated (*θ*_*E*_ ) states. **c** A box-plot of angles of microtubules (Contracted: N = 4 cells, 240 measurements. Elongated: N = 3 cells, 200 measurements). As the cell relaxes and elongates, microtubules uncoil and therefore, the angle made with the transverse axis of the cell increases. Gray lines in the middle of the boxes represent the median of the data set; the edges of the box represent the quartiles where the bottoms represent 25% and the tops represent 75% percentiles. The whiskers extend to the most extreme non-outlier data points. **d** A physical model of the microtubules. Only one helical wire is used to resemble a single microtubule, as the wire rotates to contract, the angle increases.

Thus, the antagonistic roles of myoneme and microtubules is evident. The myonemes generate contractile forces to power the rapid contraction phase, leading to bending of the transversely aligned microtubule arrays into a higher energy state (like a spring). This stored bending energy powers the relatively slow elongation back to the original cell length. We provide support for this microtubule bending model by using a helically coiled metal spring, which represents the transversely aligned microtubules on the cell body (Fig. 3d, SI Movie 4). It can be readily seen that a manually pushed spring closer (small *θ*) will spontaneously extend to relieve the stored spring energy and extend to the original length (large *θ*).

### Buckling of the Plasma Membrane

How does the plasma membrane deform during contraction? The cell membrane presents a physical barrier between the cell and the surrounding environment. Since the membrane also integrates and adheres to the cytoskeletal components underneath [35], we anticipate that during Spirostomum’s full body contraction, the incompressibility (surface area is conserved) of the membrane will lead to dramatic bucklings. Using a cell membrane stain, we observe that indeed during the contraction the membrane significantly buckles as shown in Fig 4a,b. We quantify this buckling by defining a membrane packing factor *pf* = *L_m_/L*, where *L_m_* is the membrane contour length and *L* is the projected length in the x-direction (Fig. 4c). We find that the membrane packing factor doubles from *pf* = 1.0±0.2 to *pf* = 2.1±0.2, revealing the extreme deformability of the membrane during contraction (Fig 4d). Physically, we hypothesize that the out-of-plane buckling is similar to features observed on compression of any incompressible layer, such as the skin. For repeated contraction-extraction cycles, we hypothesize that this robust elastic membrane serves to keep intracellular organelles and cytoplasmic fluid from bursting out and may serve as a useful system to investigate the role of membrane deformation in mechanosensing in living systems [36].

**FIG. 4.**
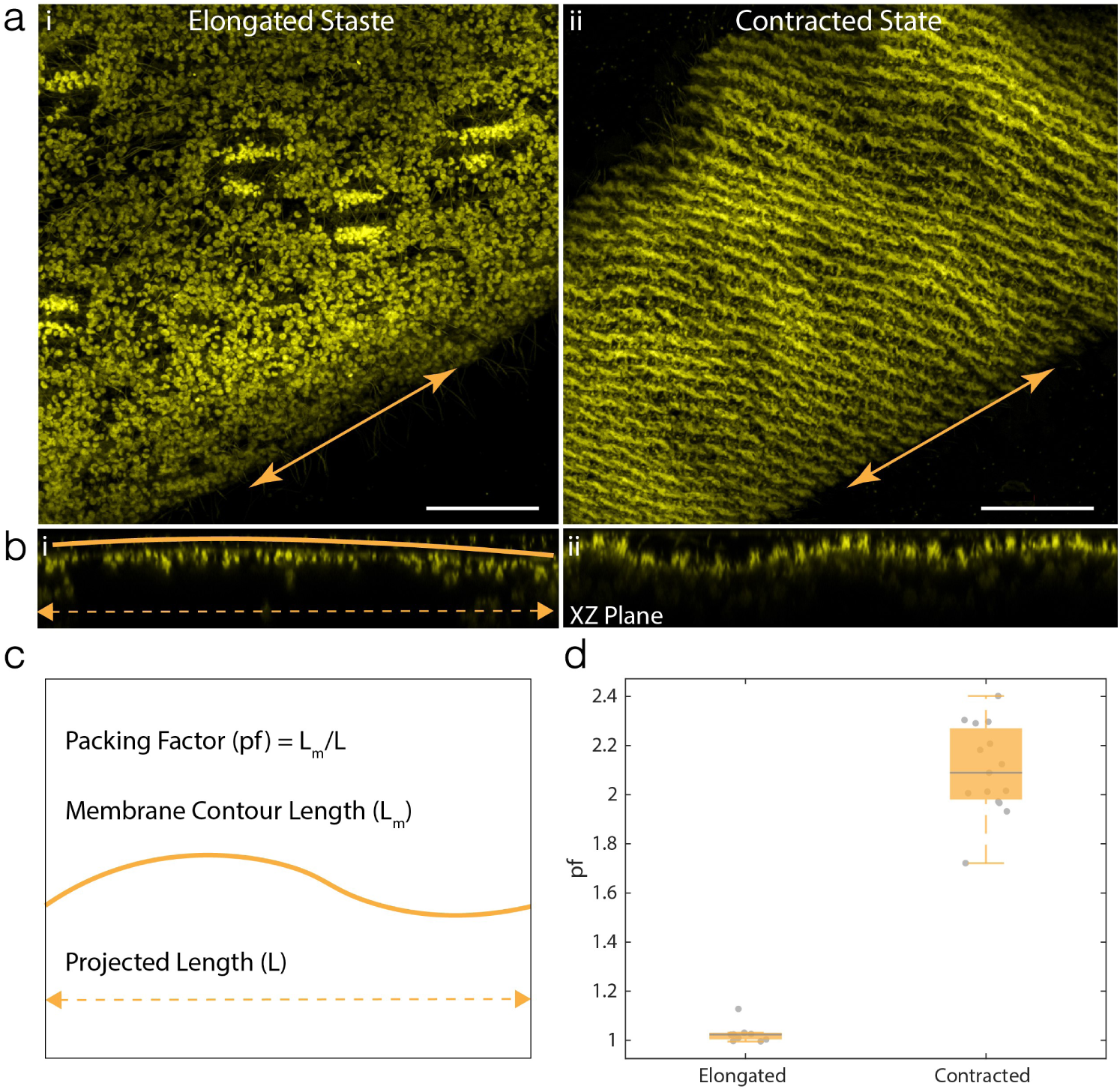
Buckling of the Plasma Membrane. **a** Immunofluorescence labelling of the cell membrane in both (i) elongated and (ii) contracted states. Orange double arrows represent the longitudinal axis of the cell. Scale Bar = 20 *µ*m. **b** XZ plane of the 3D data, with the orange curve indicating the membrane contour, and the double-arrow indicating the projected length in the x-direction. For elongation state, cell membrane is very smooth, whereas for contraction state, cell membrane is buckled. The XZ plane is sliced along the longitudinal axis of the cell. **c** Definition of the Packing Factor (*pf*) **d** A box plot showing the extent of membrane packing before and after the contraction. As the cell contracts, cell membrane buckles (Contracted: N = 2 cells, 10 measurements. Elongated: N = 3 cells, 15 measurements).

### A Physical Model to Explain Same Direction Twisting Contraction

Putting together the physical analogs for both the myonemes and the microtubules, we can now address the puzzling twisting observation during contraction (Fig 1b, SI Movie 2). When Spirostomum contracts, the cell undergoes twisting on both ends in the same direction (with a small delay), which leads to shortening of the cell body length. Applying these two simple observations of i) same-twist rotation and ii) shortening of cell body length to the physical model (Fig 5, SI Movie 5), the physical model qualitatively captures the observed Spirostomum contraction behavior. Indeed from this physical model it becomes quite clear that the rotation of both ends in the same direction is the only way to ensure bulging of the cell body to conserve cell volume and shortening of the cell body at the same time. If the rotation on the two ends are in opposite directions the cross-sectional diameter decreases and prevents the cell from shortening in length. Alternatively, since both ends of cell are free (untethered), when one end rotates, it builds up torsional stress, which is relieved by the other end rotating in the same direction.

**FIG. 5.**
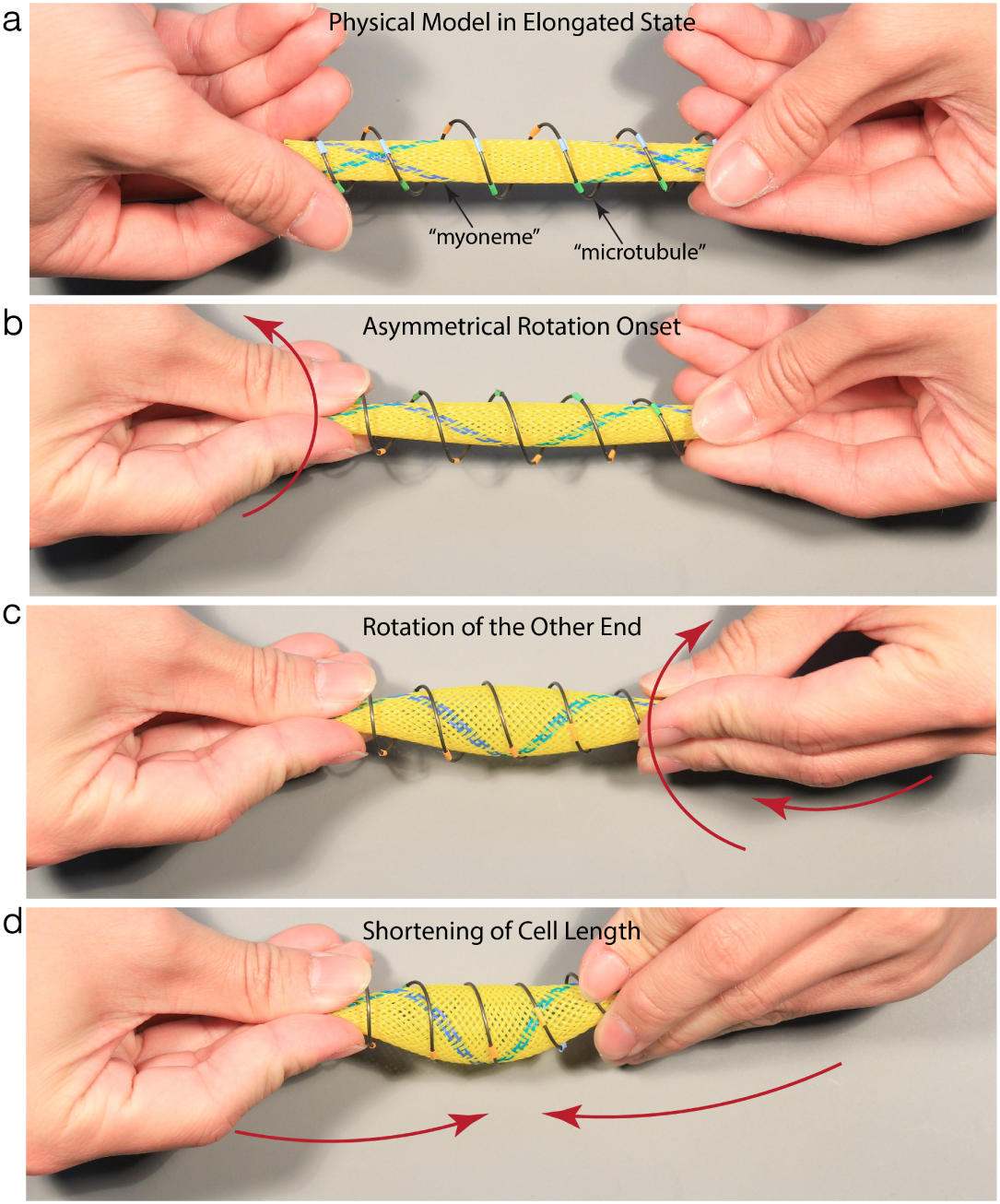
A Physical Model of the Contraction Mechanism. **a** A model of an elongated Spirostomum. The metal helical spring represents a single microtubule and the yellow mesh represents the myoneme. The wire and the mesh are connected at the ends to avoid slipping past each other. **b** As observed from high-speed data (SI Movie 2), contraction initiates at one end, and then propagates along the length of the cell. As the cell contracts, the microtubules get compressed (decreased *θ*), loading up bending energy in their deformation (loading up of a helical spring). **c** Due to the rotation that initiates from the first end, the microtubules rotate in the same direction to relieve any torsional stress, and achieve only a pure compression. Therefore, the other end has to rotate in the same direction. **d** Finally as the model is pushed closer, the myoneme parallelograms shrink and the microtubule angles decrease, to contract the entire cell with a bulge in middle region (to keep the effective volume constant).

## Conclusions

For the first time, we show that the myonemes, micro-tubules, and plasma membrane in the entire cell in both the elongated and contracted state. We first quantify the individual myoneme parallelograms, the angle change in microtubules, and the buckles in the plasma membrane of Spirostomum. Additionally, we have built a physical model that accurately captures the same-direction twisting of the two ends. Although we have presented the model to capture the contraction mechanism, open question remains about the exact integration of myoneme, microtubules, and the plasma membrane during the contraction. Answers to this question can advance our understanding of extreme motion in unicellular organisms and also inspire design of synthetic actuators.

## ACKNOWLEDGEMENTS

We thank all members of the Bhamla Lab for their feedback, and the Georgia Tech Microscopy Core for providing training on confocal microscopes. M.S.B acknowledges funding support through NSF (award no. 1817334)

## AUTHOR CONTRIBUTIONS

M.S.B. and L.X.X. designed and improved the experimental protocol, L.X.X. wrote the analysis codes and performed all characterization and analysis. Both authors contributed to writing of the manuscript.

